# Non-invasive sex genotyping of paiche *Arapaima gigas* by qPCR: An applied bioinformatic approach to identify sex differences

**DOI:** 10.1101/2021.02.12.430980

**Authors:** Edgar A. López-Landavery, Guillermo A. Corona-Herrera, Luis E. Santos-Rojas, Nadhia M. Herrera-Castillo, Tomás H. Delgadin, Sandra Tapia-Morales, Sophia González-Martinez, Lorenzo E. Reyes-Flores, Alan Marín, Carmen G. Yzásiga-Barrera, Juan I. Fernandino, Eliana Zelada-Mázmela

## Abstract

*Arapaima gigas*, one of the largest freshwater fish in the world, is suffering from high fishing pressure and habitat loss, which threaten the conservation status of its natural populations. Of great cultural importance to Amazonian people, the paiche or pirarucu *A. gigas* is in high demand for its meat, ornamental uses and other byproducts such as scales. Aquaculture is a feasible solution to this dilemma. However, the fact that *A. gigas* presents no sexual dimorphism until it is 5 years old and its long period to sexual maturity are major obstacles for brood-stock management and fingerling production. Thus, the aim of this study was to develop a molecular tool for non-invasive genotypic sexing of paiche throughout its life cycle. We collected samples from gonads, fins and gill mucus of juvenile specimens from local facilities for histological and molecular analysis. Based on the recently available genome sequence of the paiche and making use of current NGS method, we implemented a novel approach, called Genome Differences by Unmapped Reads, to identify DNA sex markers. We found a Male-Specific Region (MSR), identified as MSR_3728, to be present only in males. Next, we designed two specific sets of primers on this region to identify genotypic sex by qPCR assays. Both primer sets, MSR_107 and MSR_129, detected males with 100% accuracy. Then we developed a duplex qPCR reaction for each primer set along a reference gene, analyzed the melting curves and detected males by observing two distinct peaks, one for MSR_107 or MSR_129 and one for the reference, while females only presented the reference peak. The same results were obtained for gonads, fins and interestingly, a non-invasive source from gill mucus samples. Finally, the gonads were evaluated histologically in a double-blind test, showing 100% accuracy with qPCR assay for identifying males and females. Data clearly demonstrated a novel pipeline approach for identifying DNA sex markers, followed by a quick, non-invasive, cost-effective duplex qPCR method for sexing A. *gigas*. These results may be valuable to efficient paiche aquaculture and conservation studies, helping to reduce the fishing pressure on its natural populations.

**Highlights of the manuscript:**

– Implementation of a novel approach, called Genome Differences by Unmapped Reads, to identify DNA sex markers.
– Finding of a Male-Specific Region (MSR), present only in males.
– Development of a duplex qPCR to identify genotypic sex through non-invasive sampling.

## Introduction

*Arapaima gigas*, also known as paiche in Peru and pirarucu in Brazil, is one of the largest freshwater fish species in the world. It inhabits floodplains, rivers and lagoons in the Amazon basin (Nelson et al., 2016; Carvalho et al., 2017). Its natural distribution encompasses the South American countries Brazil, Colombia, Ecuador, Guyana, Peru, Suriname, and Venezuela (Chu-Koo et al., 2008; Hrbek et al., 2005; Farias et al., 2019, de Oliverira et al., 2019), and it was introduced to the Bolivian Amazon in the late 1970s (Miranda-Chumacero et al., 2012). Owing to its high value as an ornamental fish species, it can also be found in Southeastern Asian countries (e.g., Indonesia; Marková et al., 2020). The paiche has great cultural and historical importance for the Amazon region. It has been used as a meat supply for local people for decades, and as consequence of high fishing pressure and habitat loss, its natural population has been declining (Hrbek et al., 2005; Castello and Stewart, 2010; Castello et al., 2015; Farias et al., 2019). Importantly, A. gigas has been listed in Appendix II of CITES for protection since 1975. However, the declining population size of *A. gigas* contrasts with the high demand for its meat and other byproducts from local and potential new foreign consumers. In this conflictive scenario, *A. giga*s aquaculture appears as an ideal solution, which may benefit its conservation status, consumer demand and local producers.

In addition to the biological importance of *A. gigas* from an ecological and conservation standpoint, in recent decades it has been used for aquaculture due to its striking features (Saint-Paul, 1986; Pereira-Filho, 2003). It has one of the highest growth rates in fish, easily reaching a body weight of 10 kg during the first year of life and as much as 200kg as an adult (Saint-Paul, 1986; de Oliveira et al., 2012). Moreover, it is an obligate air-breathing fish with a large, highly vascularized swim-bladder that functions as an air-breathing organ and enables it to inhabit the hypoxic floodplain waters in the Amazon basin, which would facilitate rearing individuals at high densities in ponds (Pereira-Filho, 2003; de Oliveira et al., 2012; Val and Almeida-Val, 2012). Nevertheless, not much is known about its reproductive biology.

*A. gigas* attains sexual maturity at 5 years of age (Fontenele, 1948; Guerra Flores, 1980; Sandoval, 2007), when males and females may display different body color patterns (Saavedra Rojas et al., 2005). After a pair is formed, the female prepares a nest in sandy soils to spawn the eggs, which are immediately fertilized by the male. Both male and female adults display parental care during the first days after hatching, secreting a characteristic mucus from the “secretory organ” located at the top of the head, which is thought to provide nutrients to the fries, although only the male continues this behavior until fingerlings are left free (Noakes, 1979; Sandoval, 2007; Torati et al., 2017).

Other than color pattern during the breeding period, A. gigas presents no evident sexual dimorphism. The lack of a quick, reliable means to identify its sex before sexual maturation hampers the breeding management and fingerling production involved in rearing it in captivity. Although there have been recent efforts to resolve this situation (Chu-Koo et al., 2008; Torati et al., 2017), one of which resulted in an immune assay method for vitellogenin and sexual steroid determination, there is still a need for a quick method. Taking advantage of the recent availability of the *A. gigas* genome sequence and based on newly-developed bioinformatic methods for determining sexual markers, we aimed to develop a duplex qPCR method for a rapid, non-invasive, robust genotypic sex identification of *A. gigas* at any developmental stage.

## Materials and Methods

### Experimental organisms, biometrics and biological samples

Juvenile (18-month-old) *A. gigas* males (n=27) and females (n=28) were obtained from aquaculture farms in the city of Pucallpa, department of Ucayali, Peru. Their total length and mass, and gonad length and mass, were measured to calculate condition factor (Htun-Han, 1978) and gonadosomatic index (Anderson et al., 1983). Fish were handled in accordance with the Universities Federation for Animal Welfare Handbook on the Care and Management of Laboratory Animals (www.ufaw.org.uk) and internal institutional regulations.

For molecular and histological identification, gonads (from 28 females and 27 males) and pectoral fin clip (16 females and 13 males) samples were taken from each specimen and preserved in 100% ethanol until molecular analysis. Additionally, gonad samples were taken and preserved in 10% neutral buffered formalin (NBF 10%) for phenotypic sex identification by histological procedures. Finally, gill mucus (16 females and 13 males) was sampled with sterile drugstore cotton swabs (Cody et al., 2018) and preserved in DNA lysis buffer solution until molecular analysis.

### Gonad histology

The gonad samples, fixed in NBF 10%, were rinsed and transferred to 70% alcohol, and processed in the automatic dehydrating and paraffin inclusion device Thermo Scientific™ STP 120 Spin Tissue Processor. The tissue was embedded in a modular Tissue Embedding Center EC 350, and 6 µm sections were prepared with a Rotary microtome/ Semi-automatic CUT 5062 SLEE Medical and stained with Hematoxylin-Eosin standard protocol in the Thermo Scientific Varistain® V24-4 AUTOMATIC SLIDE STAINER. The histological slides were mounted and observed under a Motic BA310 Compound Microscope and microphotographs were taken with a Moticam 10.0 MP.

### De Novo genome assembly

Illumina NGS sequences (250 bp) were obtained from *A. gigas* females (n = 2) and males (n = 2) (Du et al., 2019), hosted in the database SRA (SRX7250390, SRX7250391) from NCBI. As a first step, the reads were analyzed with the FASTQC program (Andrews, 2010) to determine whether they met the minimum quality necessary to carry out the assembly.

Subsequently, the low-quality reads (Q < 28), adapters and “N” bases were removed with the Trimmomatic program (Bolger et al., 2014). Then duplicates were eliminated with the Deduped program (Gregg and Eder, 2019)) and corrected with the SOAPdenovo Corrector or SOAPec program (Li et al., 2010). Next, a second FASTQC quality control analysis (Andrews, 2010) and a results report were performed with the MultiQC software (Ewels et al., 2016) to corroborate the quality of the reads before the assembly process. Subsequently, the genomic assembly was carried out with the Discovardenovo program (Weisenfeld et al., 2014). Finally, the male and female genomic assembly quality was evaluated with BUSCO (Simão et al., 2015), Genome Scope (Vurture et al., 2017), Quast (Gurevich et al., 2013) and Bowtie2 (Langmead and Salzberg, 2012).

### Male-Specific Region (MSRs) identification by the Genome Differences by Unmapped Reads (GDUR) pipeline

After the male and female genome assemblies were obtained, a search was performed for sex-specific candidate regions. For this purpose, we implemented a bioinformatic pipeline, which we called “Genome Differences by Unmapped Reads (GDUR)”. It is based on multiple programs that are currently used for mapping and filtering NGS sequences, but which have not been used together to obtain sex-specific sequences.

As a first step, we mapped independently the forward and reverse male reads vs. the female *A. gigas* genome with the Fastq Screen program (Wingett and Andrews, 2018). We focused on the unmapped male reads because they have a greater probability of being male-specific sequences. Subsequently, the pair sequences (Forward and Reverse) of the unmapped male reads were searched using the Fastq Pair program (Edwards and Edwards, 2019). Once the sequences with their corresponding pairs were obtained, the cleaning and preparation steps of the reads were performed with the Trimmomatic (Bolger et al., 2014), Deduped (Gregg and Eder, 2019), SOAPec (Li et al., 2010), FastQC (Andrews, 2010) and MultiQC (Ewels et al., 2016) programs. After that, the genomic sex sequences of the *A. gigas* male were assembled with Discovardenovo and evaluated with BUSCO, Genome Scope, Quast and Bowtie2, as explained in the previous section. Then, local BLAST (Altschul et al., 1990) alignments of the male-specific sequences vs. the female genomic assembly were carried out to corroborate that male sex-specific sequences are not present in the female genome.

Similarly, a local BLAST (Altschul et al., 1990) of the sex-specific sequences vs. the male genomic assembly was carried out to corroborate that the sequences are found in the genome assembly of *A. gigas* males. In addition, with some of the identified sequences, forward and reverse pairs (Fw-Rv) of Male-Specific Region (MSR) and reference gene primers were designed with Primerquest and the oligo analyzer tools of IDT (www.idtdna.com). The designed primer sets were used to perform experimental validation by PCR and qPCR analysis.

### DNA extraction and qPCR experimental validation of Male-Specific Regions (MSRs)

*A. gigas* male and female DNA was extracted from gonads, fins and mucus by the standard phenol-chloroform method added with proteinase K and RNase (Mirimin et al., 2015; Dominguez et al., 2018). The integrity of DNA was observed on a 1% agarose gel electrophoresis with TBE 0.5x and 100 V for 60 min (Simbolo et al., 2013). The purity (A260/ A280) and concentration (ng/µl) values were obtained with an Epoch Microplate Spectrophotometer (www.biotek.com) and a Qubit fluorometer® 3.0 using a QubitTM dsDNA BR assay kit (https://www.thermofisher.com/), respectively. Preliminary PCRs were carried out with DNA from male and female gonads with a PCR Master Mix 2x (https://www.thermofisher.com/) and a male-specific or reference gene final primer concentration of 0.5 µM and 100 ng of total DNA. Under a cycling protocol of initial denaturation step 5 min at 95°C, final extension 5 min at 72oC and 35 cycles of amplification (Denaturation 94oC, Annealing 60oC, Extension 72oC x 20 s) in a Veriti 96-Well Thermal Cycler (https://www.thermofisher.com/). Finally, the presence of one specific PCR product in males and no product in females (MSR primers), and one product in both sexes (reference gene primers) were corroborated with standard 1.5% agarose gel electrophoresis (TBE 0.5x, 100 V, 45 min).

After duplex PCR experiments were designed with the MSRs and reference gene primers, corroborating the absence of dimers and unspecific products, the best duplex primers were tested by qPCRs with the SYBR-Green master I protocol, in a Roche LightCycler 480 II real-time PCR instrument (https://lifescience.roche.com/). The qPCR reactions were optimized with 30 ng of total DNA, and a final primer concentration of 0.05 µM, with a protocol of initial denaturation step 5 min at 95°C, final extension 5 min at 72°C, 35 cycles of amplification (Denaturation 94°C, Annealing 62°C, Extension 72°C x 20 s) and a final melting curve analysis of the PCR products.

### Statistical analysis

For biometric comparisons, the statistical differences between values were determined by Student’s t-test (P ≤ 0.05) with the GraphPad prism V. 9.0 software.

## Results

### Biometrics, condition factor and gonadosomatic index (GSI) of A. gigas

Fish sampled at the aquaculture facilities did not show any differences between males and females regarding total length, total mass and condition factor (**Supplementary Table 1**). Nevertheless, there were statistical differences in gonad mass, gonad length and GSI between males and females, with females having the largest gonads (p < 0.05; **Supplementary Table 1**).

### Genome assemblies and Male-Specific Region identification by GDUR pipeline

The genomic assembly generated a total length of bases of 657.53 Mb for females and 659.05 Mb for males. The evaluation of assemblies from both sexes indicated N50 values of approximately 75,000 bp for females and 81,000 bp for males. This meant a difference between assemblies of about 6,000 bp between sexes. The L50 values were 2,112 bp for females and 1,962 bp for males (**Supplementary Table 2**).

The coverage for the genome was 42.9x for females and 42.6x for males. In addition, genome scope analyses indicated an error rate of 0.04% in the genome reads. Comparison of the two sexes showed that the assembly of males presented a greater haploid length (628.16 Mb), a greater length of repetitions (28.077 Mb), a greater length of unique sequences (600.08 Mb), and a greater number of duplications (0.74%) than did the female assembly. Additionally, males presented lower levels of heterozygosity (0.16%) than did females (0.22%, **Supplementary Table 2**). These values suggest that males have a larger genome, with more duplicated regions, and have lost heterozygosity throughout their evolution. In this case, sex-specific genomic duplications and a possible hemizygous condition could be assumed (they only have one copy of the X chromosome genes, instead of the usual two), as occurs in male mammals with XY sex determination systems.

The evaluation of genomes by mapping the reads with Bowtie2 indicated that assembly of females presents a slightly higher number of reads (64 million) than the males; however, the mapping rate for both genomes was ∼ 9 8. 8 % (**Supplementary Table 2**). On the other hand, the analysis of the presence of orthologous groups with BUSCO indicated that assemblies presented similar values in the coincidence of identified orthologous genes (86%), with about 10% fragmented and only 4.3% lost using the “vertebrata” database. Based on statistics and quality control of genome assemblies, the next step was to proceed with the analyses to obtain sex-specific genomic sequences in *A. gigas*.

Paiche sex-specific sequences were generated with the methodology called Genome Differences by Unmapped Reads or GDUR (**Figure 1A**). As the first step, the reads of the males that were not mapped into the female genome were identified. The levels of reads not mapped in the female genome were 2.33% and 5.50% for the forward and reverse sense, respectively. This condition found with the present workflow suggests that most of the sex-specific sequences are found in the antisense strand of the male *A. gigas* genome. Once sequences were identified, their corresponding pairs were obtained from the original cleaned files. Quality control of male sex-specific paired reads generated 897,263 unique reads (91.5%) and 83,571 duplicated reads (8.5%), with average Phred quality 38, 45% GC, and average length 176 bp (**Supplementary Table 3**).

**Figure 1.**
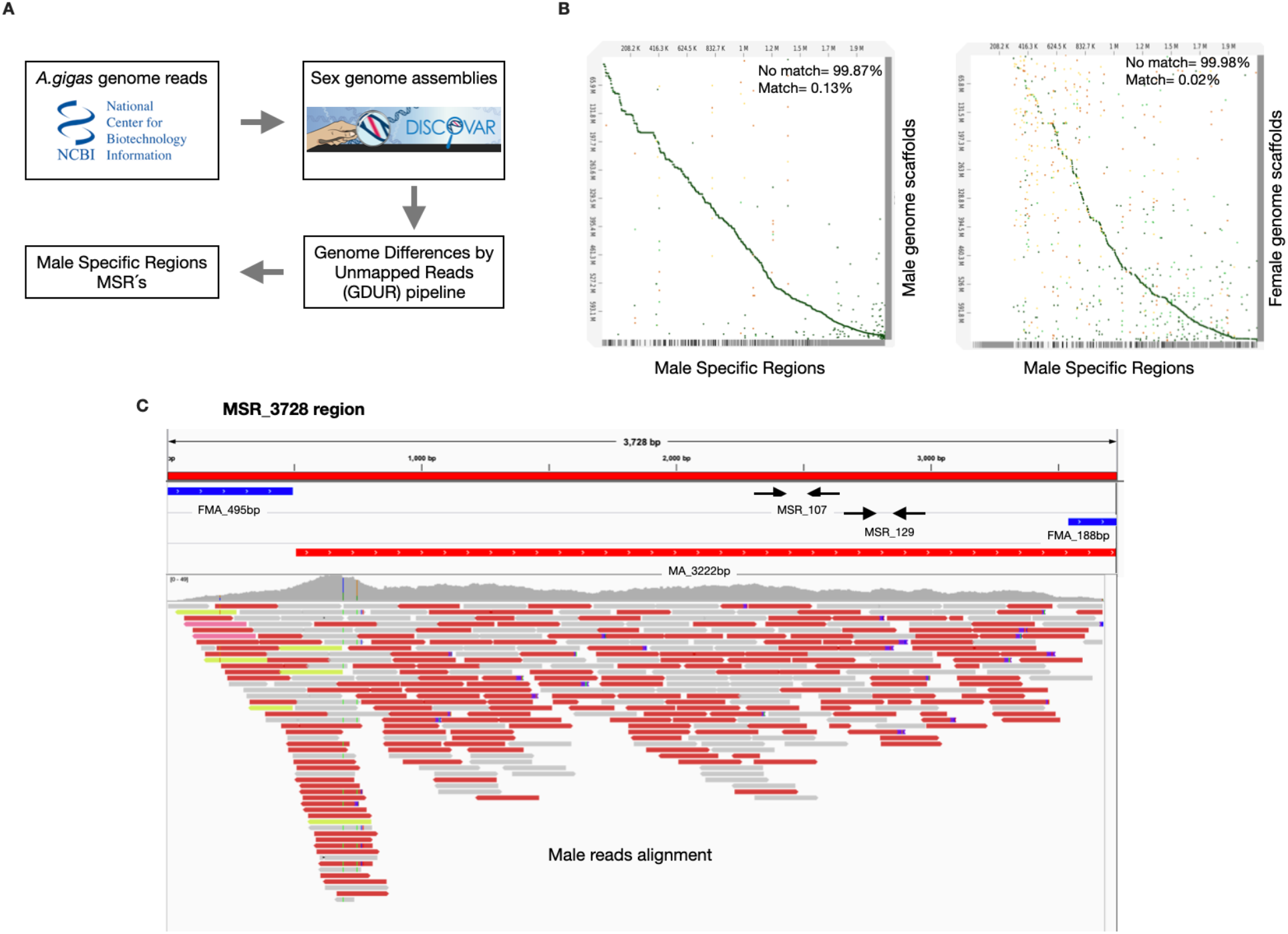
A) Summary of bioinformatic methods for finding male-specific regions in *Arapaima gigas*; B) Dot-plot alignment of male-specific regions vs. male and female genome scaffolds of *A. gigas*; C) Male-specific region (MSR_3728) used to validate the methodology by qPCR vs. Female genome alignment. (FMA) Female genome scaffolds alignment; (MA) Male genome scaffolds alignment.

The assembly of sex-specific paired reads generated an N50 value for scaffolds of 3,542 bp, with a total length of approximately 2.08 Mb, a low percentage of chimeras (0.02%) and a coverage of 93.1x. The total number of contigs for this assembly was 903, with a N50 value of 3,417 bp, 43.59% GC and with the presence of 9.82 “Ns” per 100 Kb. The analysis with BUSCO showed that only 0.4% of the orthologous genes in the vertebrata database were found in the sex-specific assembly of *A. gigas* (**Supplementary Table 3**).

The genome scope analysis indicated that the read error rate for this assembly was 1.67%, with a haploid length of 3.09 Mb, a repeat length of 102.20 bp and unique sequences length of 2.98 Mb. Furthermore, the duplication level was 1.23% and heterozygosity was 1.74%, with a model fit of 97%. Another characteristic was that the percentage of female reads that unmapped in the male sex-specific assembly was 98, suggesting that just 2% of the female reads showed some similarity with the male sex-specific assembly (**Supplementary Table 3**).

Finally, an analysis to compare the assembled genomes was performed with the D-Genies software (Cabanettes and Klopp, 2018). Comparison between MSRs and male genome indicated that just 0.13% of the sequences matched between scaffolds (**Figure 1B**). In contrast, comparison between MSRs vs. female genome clearly showed a match of just 0.02% between scaffolds **(Figure 1B)**. These results suggest very low similarity of the MSRs vs. the female genome, and demonstrate the confidence of the GDUR pipeline implemented in this study to find sex-specific sequences of *A. gigas* males.

### q PCR Primers design and experimental validation of Male-Specific Regions in A. gigas gonads, fins and mucus

After identifying the MSR_3728 region, two sets of primers on this sequence were designed to be used in the qPCR sexing tool. The first step was to check whether there were female scaffolds that matched the MSR_3728 region, obtaining the FMA_495bp at the beginning and FMA_188bp at the end of this region. Next, the MSR_3728 region was aligned against the male genome, obtaining a 3,222bp region, and corroborating its presence in males and its absence in females. Then, and based on a 3,045bp sequence in the MSR_3728 region, two pairs of qPCR primers were designed to validate the GDUR pipeline in *A. gigas* (**Figure 1C**): MSR_107 (from 2,457bp to 2,564bp) and MSR_129 (from 2,540bp to 2,669bp) (**Figure 1C**). As the present study aimed to develop a duplex qPCR controlling for DNA integrity, four reference genes (HKGs) were tested: TATA-Box Binding Protein Like 1 (TBPL1), 60S Ribosomal Protein L3 (RPL3), Eukaryotic Translation Elongation Factor 1 Alpha 1 (EF1A1), and Phosphoglycerate Kinase 1 (PGK1). Out of all the HKGs, TBPL1 was the best choice, because it produced a 205bp amplicon with no inter-or intra-specific dimeric interactions with itself or between the MSR_107 or MSR_129 primers.

For qPCR validation, DNA was extracted from female and male *A. gigas* gonads, fins and mucus (**Figure 2A**). All samples presented an A260/A280 ratio of 1.8 and concentrations between 946.5 and 1,999.6 ng/µl. Analysis of the PCR products showed two amplicons in males and only one amplicon in females, corresponding to the MSR_107 or MSR_129 product plus TBPL1_205 product in males and TBPL1_205 product in females (**Figure 2B**). Afterwards, melting curve analyses showed that the best qPCR condition was 35 amplification cycles with a primer combination of 0.1µM for MSR and 0.05µM for HKG at annealing temperature of 62°C (**Figure 2C**). PCR efficiencies obtained by the standard curve method were 109.19% and 101.32% for MSR_107 and MSR_129 primers, respectively. Efficiencies for TBPL1_205 were 96.16% for males and 98.22% for females (**Supplementary Table 4**). The qPCR melting curve analysis showed temperatures of 76.6, 77.5 and 84.7°C for the qPCR products of MSR_107, MSR_129 and TBPL1_205, respectively (**Figure 2C**). Data of melting curves and qPCR products for 14 males and 14 female gonads is shown in Figure 3, and the melting temperature and Cq values for all the samples are shown in **Supplementary Table 5**.

**Figure 2.**
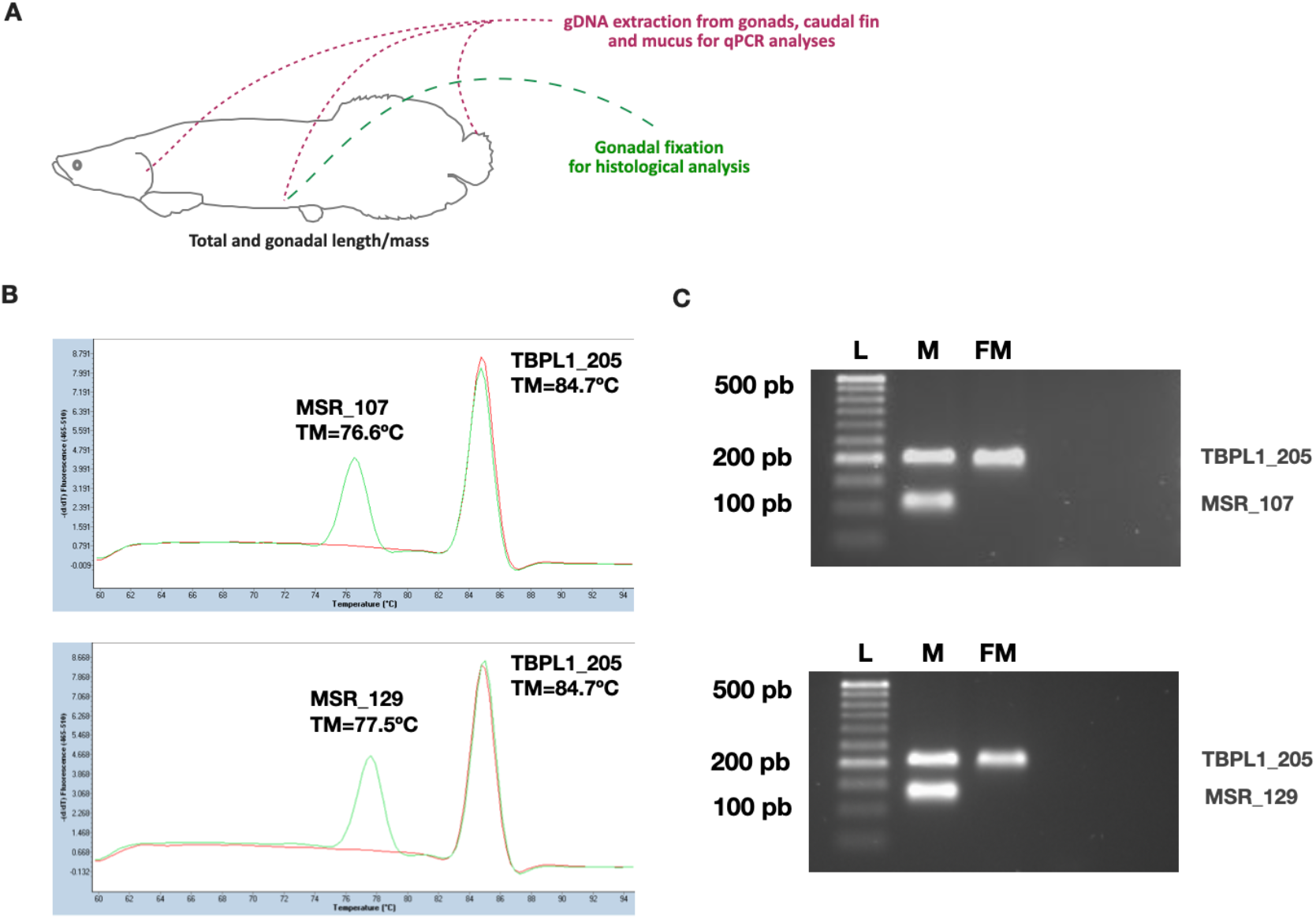
A) Summary of *Arapaima gigas* juvenile sampling; B) qPCR melting curves of Male-Specific Regions (MSR_107/MSR_129) and the reference gene TBPL1_205 in *A. gigas* males and females; C) Gel electrophoresis of duplex PCR products of Male-Specific Regions (MSR_107/MSR_129) and the TBPL1_205 gene in *A. gigas* males and females. (L) Ladder 50 bp; (M) Male; (FM) Female.

**Figure 3.**
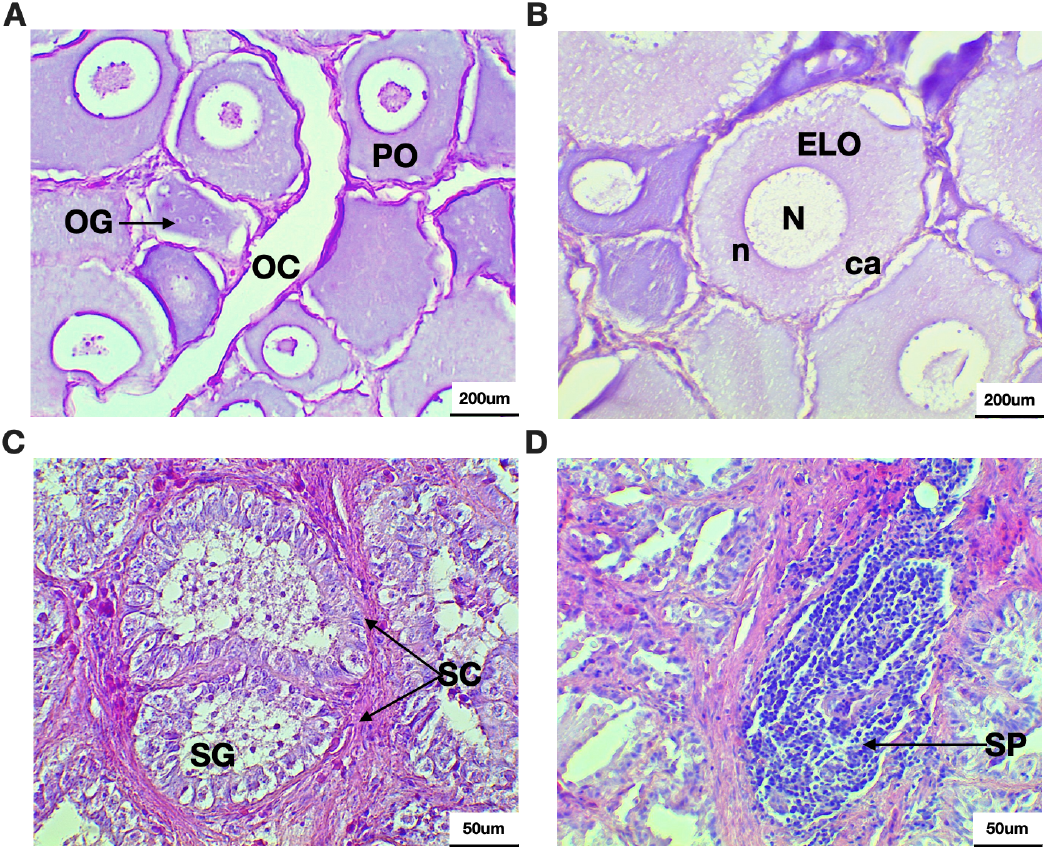
Histological observations of gonads from *Arapaima gigas* juveniles. A; Early growing ovary, B; Growing stage ovary, C; Early growing testis, D; Growing stage testis. (OC) Ovarian cavity; (OG) Oogonium; (PO) Perinucleolar oocyte; (ELO) Early lipid oocyte; N (Nucleus); (n) Nucleolus; (ca) Cortical alveoli; (SC) Spermatocytes; (SG) Spermatogonium; (SP) Spermatids.

Finally, the set of primers was tested in double-blind mode to corroborate the correlation between gonadal phenotype and genotypic sex. For 100% of the tested samples, males and females were identified at molecular level with the primers MSR_107 and MSR_129 (**Table 1**). In some mucus samples, the qPCR analysis showed a degree of DNA cross-contamination, mainly due to the nature of the samples.

**Table 1:** Percentage of sex identification with the Male-Specific Regions (MSR_107/MSR_129) in *Arapaima gigas*. (n) number of individuals.

| MSR | Source | Male | Female | Total n | Percentage of sex identification |
| --- | --- | --- | --- | --- | --- |
| MSR_107 | Gonad | 27 | 28 | 55 | 100 % |
|  | Fin | 13 | 16 | 29 | 100 % |
|  | Mucus | 13 | 16 | 29 | 100 % |
| MSR_129 | Gonad | 27 | 28 | 55 | 100 % |
|  | Fin | 13 | 16 | 29 | 100 % |
|  | Mucus | 13 | 16 | 29 | 100 % |

### Histological observations on gonads of A. gigas juveniles

Finally, gonad phenotype was identified in order to validate the double-blind performed previously with the sexual genotype. In this regard, the histological observations identified female ovaries at primary stages of development (early previtellogenesis), characterized by the presence of a clear ovarian cavity, oogonium, perinucleolar oocytes and early lipid oocytes with a nucleus, nucleolus and cortical alveoli (**Figure 3 A, B**). The testes of males were found at primary developmental stages, characterized by clear spermatocytes with spermatogonium, testicular lumen and in some cases a slightly presence of spermatids (**Figure 3 C, D**). The histological observations and the GSI data clearly indicate that 18-month-old *A. gigas* juveniles are at immature gonadal stages, as described in Yong et al. (2020), for fish at same age.

## Discussion

External sexual dimorphism is present in many fish species, but absent in many others (Nelson et al., 2016). From the standpoint of aquaculture and conservation, easy sex identification is needed to manage reproduction and measure the health of a fish population. There is great interest in the aquaculture and conservation of *A. gigas* (paiche) (Hrbek et al., 2005; Castello and Stewart, 2010; Castello et al., 2015; Farias et al., 2019), which is a very important Amazonian fish. It presents no sexual growth dimorphism, and as far as we know, there is no other published data reliable enough to identify sex throughout its life cycle. Thus, the lack of convenient methods for sex identification, which impairs breeding management and fingerling production, is a major obstacle for paiche culture. The *A. gigas* genome has recently been published (Du et al., 2019), providing the opportunity to search for sex-specific genetic markers which would enable genotypic sex to be identified at any stage in the life cycle by means of a simple method. As mentioned above, one of the main difficulties for efficient breeding management of *A. gigas* in captivity is the lack of an adequate method for identifying sex. Thus, after identifying a male-specific region (MSR) for *A. gigas*, we developed a duplex qPCR to identify genotypic sex, which provided highly accurate sex identification for all samples, with 100% match between genotypic sex (PCR) and biological sex by morphological identification of the gonad (histology). Moreover, this qPCR method was applied successfully not only to tissue samples (i.e., gonads and fins) but also to gill mucus, which makes this a truly non-invasive, novel technique for sexing fish.

The identification of genotypic sex in fish is being increasingly used thanks to the availability of NGS techniques for species for which aquaculture and conservation are important. Recently, an approach using NGS method and PCR validation was used on the mandarin fishes *Siniperca scherzeri* and *Siniperca knerii* (Han et al., 2020). These studies designed three set of primers on a MSR which identified males with a success rate of 100%. Similar PCR protocols based on sex-specific markers have been developed for other fish species, for instance in the carps *Hypophthalmichthys nobilis, Hypophthalmichthys molitrix* (Liu et al., 2018) and the snakehead Channa argus (Ou et al., 2017). However, all these methods were based on end-point PCR, which does not enable the simultaneous detection of multiple genes and requires tissue samples for DNA extraction, affecting the desired non-invasive attribute. Importantly, our PCR-based sex identification tool is a duplex Real Time PCR, where we can measure both the sex-specific marker and the HKG in the same reaction well, a desirable feature for minimizing testing costs. In fact, we were able to detect two peaks in the melting curve analysis with either combination of HKG plus the sex-specific primer set, which indicates that the duplex reaction can efficiently amplify both amplicons in every well.

Although previous work based on immunoassays for vitellogenin and sexual steroids hormones has been conducted for *A. gigas* with a high degree of success (Chu-Koo et al., 2008), it requires the handling of animals, which in the case of *A. gigas*, due to its large size, requires very laborious logistics to obtain blood samples. Moreover, it is only effective once the fish has begun its reproductive stage, which happens as from three years of age, making this tool unviable when early identification is needed for managing breeding stock. Other methods, such as endoscopies or surgery are more difficult because they are invasive, risky and require sedation, although good progress has been made in this regard (Torati et al., 2017). Importantly, however, none of these paiche sexing approaches can be applied at early developmental stages or juvenile/ resting state. Our innovative non-invasive qPCR method enables early sex identification in *Arapaim*a and can be used for studying biological traits such as growth during any developmental stage, performing nutrition tests and evaluating natural populations. However, when mucus samples are collected in rearing ponds, maximum care is required in order to minimize the chance of any possible cross-contamination for DNA amplification. Despite this small contamination to be expected given the nature of sampling, sex identification at molecular level is still reliable, because the qPCR product and the shape of the curve/band are intense, clear and well defined. Overall, these results indicate that primers MSR_107 and MSR_129 are male-specific and can feasibly be implemented as molecular markers to differentiate *A. gigas* genotypic males and females.

On the other hand, and according to the statistical results, the quality of the *A. gigas* female and male genomic assemblies was good. Based on the above, the bioinformatics pipeline described here may be considered as preliminarily validated to perform *de novo* genomic assemblies in several species of interest, with high probability of identifying sex-specific sequences to be used as molecular markers in aquaculture and conservation activities. Another interesting aspect of the male genome statistics is that they suggest that *A. gigas* has an XY determination system, where the male has the heterogametic sex. However, further bioinformatic, molecular, and cytogenetic evidence is required to corroborate this hypothesis.

Finally, the modest percentage of reads alignment with the male-specific assembly may be explained by the natural accumulation of repetitive sequences observed in the sex-specific regions of vertebrates. With this type of sequences, it is difficult to make a complete assembly when there are no long sequencing reads that enable this type of repetitive sequence patterns to be resolved. Despite the percentage of alignment and the lack of identification of orthologous genes, preliminary identification was possible for some protein sequences which may be good candidates for use as sexual molecular markers in *A. gigas*.

## Conclusion

The present work has developed a qPCR method for quick, accurate sex determination of A. gigas individuals at any stage of development. It was validated for gonad, fin and mucus samples, enabling non-invasive testing, with 100% of fish correctly sexed. This qPCR method is of great value to aquaculture and would be an excellent tool for ecological studies of threatened wild populations of *A. gigas*, as only a sample of mucus or fin suffices to sex the fish accurately.

### CRediT author statement

EALL, GACH: Conceptualization, Software, Methodology, Formal Analysis; LESR, NMHC, STM, SGM, LERF, AM, CGYB: Methodology; THD: Writing-Original draft preparation, JIF: Funding acquisition, Conceptualization, Investigation, Supervision; Writing-Original draft preparation, Writing-Reviewing and Editing; EZM: Funding acquisition, Conceptualization, Investigation, Supervision.

## Supplementary Files

**Supplementary Figure 1: A, B)** Gel electrophoresis of duplex PCR products of Male-Specific Regions (MSR_107/MSR_129) and the TBPL1_205 gene in *Arapaima gigas* males; C, D) Gel electrophoresis of duplex PCR products of Male-Specific Regions (MSR_107/MSR_129) and the TBPL1_205 gene in *Arapaima gigas* females; E, F) qPCR melting curves of Male-Specific Regions (MSR_107/MSR_129) and the reference gene TBPL1_205 in *Arapaima gigas* males and females.

**Supplementary Table 1:** Fish total weight, total length, condition factor, gonad weight and length, and Gonadosomatic Index (GSI) of *Arapaima gigas* analyzed in this study (NA: Not available).

**Supplementary Table 2:** Statistical summary for the male and female genome assemblies and assessment of their quality in *Arapaima gigas*.

**Supplementary Table 3:** Statistical summary of assembly from male-specific reads in *Arapaima gigas*.

**Supplementary Table 4:** Genes of interest(1) and housekeeping genes(2), efficiency (E) and R2 values from duplex qPCR assays.

**Supplementary Table 5:** Raw data of duplex qPCR assays for each individual tested in this study (UDT: Undetected; NA: Not available).

## Acknowledgments

The authors gratefully acknowledge the support from Universidad Nacional del Santa (UNS).

