## Supplementary files for "Non-invasive sex genotyping of paiche *Arapaima gigas* by qPCR: An applied bioinformatic approach to identify sex differences"

**Supplementary Figure 1:** A, B) Gel electrophoresis of duplex PCR products of Male-Specific Regions (MSR_107/MSR_129) and the TBPL1_205 gene in *Arapaima gigas* males; C, D) Gel electrophoresis of duplex PCR products of Male-Specific Regions (MSR_107/MSR_129) and the TBPL1_205 gene in *Arapaima gigas* females; E, F) qPCR melting curves of Male-Specific Regions (MSR_107/MSR_129) and the reference gene TBPL1_205 in *Arapaima gigas* males and females.

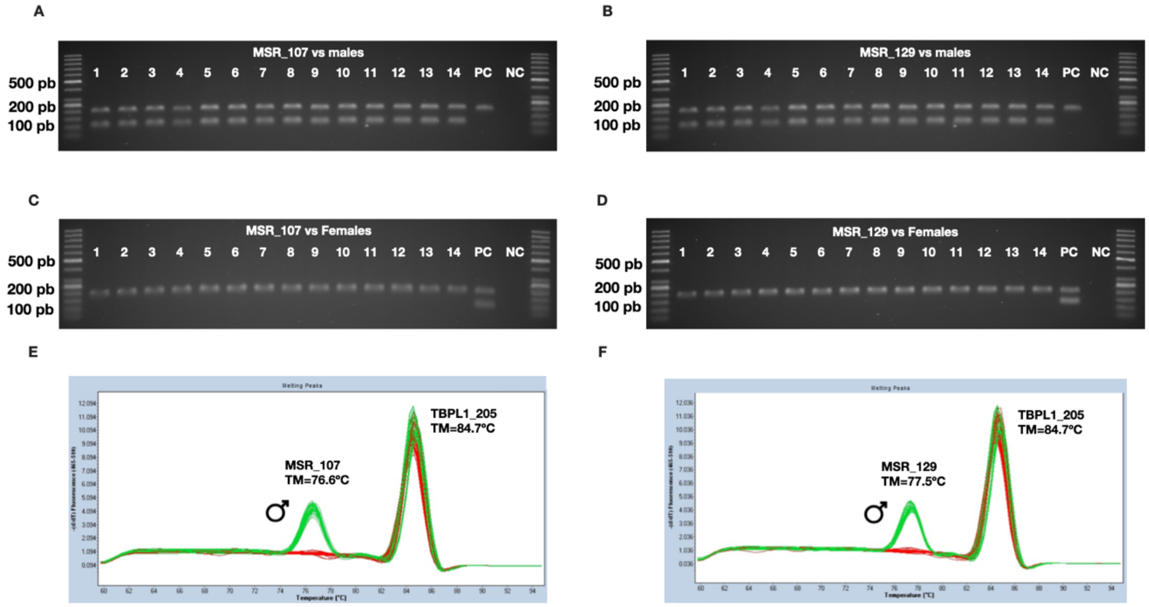

**Supplementary Table 1**. Fish total weight, total length, condition factor, gonad weight and length, and Gonadosomatic Index (GSI) of *Arapaima gigas* analyzed in this study (NA: Not available).

| ID | Total weight (kg) | Length (cm) | Condition factor | Gonad | | GSI |
|---|---|---|---|---|---|---|
| | | | | Weight (g) | Length (cm) | |
| Female_1 | NA | NA | NA | NA | NA | NA |
| Female_2 | NA | NA | NA | NA | NA | NA |
| Female_3 | NA | NA | NA | NA | NA | NA |
| Female_4 | 14.60 | 117 | 0.91 | 6.50 | 21.00 | 0.045 |
| Female_5 | 17.50 | 129 | 0.82 | 6.70 | 22.00 | 0.038 |
| Female_6 | 15.85 | 121 | 0.89 | 5.60 | 18.00 | 0.035 |
| Female_7 | 20.55 | 128 | 0.98 | 3.80 | 22.00 | 0.018 |
| Female_8 | 18.80 | 129 | 0.88 | 7.20 | 22.00 | 0.038 |
| Female_9 | 16.10 | 126 | 0.80 | 7.10 | 23.00 | 0.044 |
| Female_10 | 14.50 | 120 | 0.84 | 7.80 | 21.00 | 0.054 |
| Female_11 | 15.10 | 124 | 0.79 | 6.70 | 23.50 | 0.044 |

| | | | | | | |
|---|---|---|---|---|---|---|
| Female_12 | 14.05 | 120 | 0.81 | 6.60 | 21.00 | 0.047 |
| Female_13 | 14.20 | 117 | 0.89 | 8.60 | 23.00 | 0.061 |
| Female_14 | 17.00 | 125 | 0.87 | 9.90 | 22.00 | 0.058 |
| Female_15 | 18.75 | 136 | 0.75 | 11.50 | 28.50 | 0.061 |
| Female_16 | 15.30 | 123 | 0.82 | 10.30 | 24.50 | 0.067 |
| Female_17 | 15.30 | 121 | 0.86 | 9.20 | 19.00 | 0.060 |
| Female_18 | 12.25 | 113 | 0.85 | 9.60 | 22.00 | 0.078 |
| Female_19 | 14.75 | 118 | 0.90 | 6.70 | 21.00 | 0.045 |
| Female_20 | 16.90 | 126 | 0.84 | 9.90 | 23.00 | 0.059 |
| Female_21 | 16.85 | 125 | 0.86 | 6.50 | 22.00 | 0.039 |
| Female_22 | 17.65 | 128 | 0.84 | 7.00 | 21.00 | 0.040 |
| Female_23 | 15.65 | 133 | 0.67 | 10.20 | 22.00 | 0.065 |
| Female_24 | 20.25 | 135 | 0.82 | 7.00 | 23.00 | 0.035 |
| Female_25 | 19.05 | 127 | 0.93 | 6.50 | 23.00 | 0.034 |
| Female_26 | 15.10 | 119 | 0.90 | 6.10 | 20.00 | 0.040 |
| Female_27 | 14.70 | 128 | 0.70 | 7.20 | 23.00 | 0.049 |
| Female_28 | 14.85 | 132 | 0.65 | 6.70 | 22.00 | 0.045 |
| *Mean* | *16.22* | *124.80* | *0.83* | *7.64* | *22.10* | *0.048* |
| *SD* | *2.07* | *5.92* | *0.08* | *1.79* | *1.94* | *0.013* |

| | | | | | | |
|---|---|---|---|---|---|---|
| Male_1 | NA | NA | NA | NA | NA | NA |
| Male_2 | NA | NA | NA | NA | NA | NA |
| Male_3 | NA | NA | NA | NA | NA | NA |
| Male_4 | 17.90 | 143 | 0.61 | 3.30 | 23.00 | 0.018 |
| Male_5 | 14.35 | 112 | 1.02 | 1.80 | 16.00 | 0.013 |
| Male_6 | 16.15 | 118 | 0.98 | 1.50 | 23.00 | 0.009 |
| Male_7 | 19.70 | 129 | 0.92 | 1.50 | 16.00 | 0.008 |
| Male_8 | 19.30 | 130 | 0.88 | 1.20 | 15.00 | 0.006 |
| Male_9 | 15.60 | 123 | 0.84 | 1.10 | 16.00 | 0.007 |
| Male_10 | 18.05 | 123 | 0.97 | 1.50 | 20.00 | 0.008 |
| Male_11 | 19.10 | 127 | 0.93 | 1.40 | 19.00 | 0.007 |
| Male_12 | 16.85 | 128 | 0.80 | 1.40 | 21.00 | 0.008 |
| Male_13 | 16.05 | 126 | 0.80 | 3.20 | 20.00 | 0.020 |
| Male_14 | 16.65 | 126 | 0.83 | 2.80 | 21.00 | 0.017 |
| Male_15 | 15.50 | 123 | 0.83 | 1.80 | 20.00 | 0.012 |
| Male_16 | 16.95 | 126 | 0.85 | 3.00 | 23.00 | 0.018 |
| Male_17 | 16.15 | 127 | 0.79 | 5.60 | 27.00 | 0.035 |
| Male_18 | 16.05 | 123 | 0.86 | 1.90 | 24.00 | 0.012 |
| Male_19 | 15.06 | 124 | 0.79 | 2.00 | 18.00 | 0.013 |
| Male_20 | 16.30 | 122 | 0.90 | 2.10 | 18.00 | 0.013 |
| Male_21 | 13.55 | 117 | 0.85 | 1.70 | 23.00 | 0.013 |
| Male_22 | 21.70 | 134 | 0.90 | 2.60 | 21.00 | 0.012 |
| Male_23 | 18.60 | 127 | 0.91 | 2.00 | 22.00 | 0.011 |
| Male_24 | 18.05 | 130 | 0.82 | 1.60 | 19.00 | 0.009 |
| Male_25 | 15.25 | 120 | 0.88 | 1.40 | 22.00 | 0.009 |
| Male_26 | 16.50 | 127 | 0.81 | 1.50 | 15.00 | 0.009 |
| Male_27 | 15.85 | 131 | 0.71 | 2.10 | 20.00 | 0.013 |
| *Mean* | *16.88* | *125.67* | *0.85* | *2.08* | *20.08* | *0.012* |
| *SD* | *1.86* | *6.11* | *0.09* | *0.97* | *3.09* | *0.006* |

**Supplementary Table 2.** Statistical summary for the male and female genome assemblies and assessment of their quality in *Arapaima gigas.*

| Assembly features | Male | Female |
|---|---|---|
| Total length of bases (Mb) | 659.05 | 657.53 |
| Scaffold N50 (kb) | 81 | 75 |
| Scaffold L50 (bp) | 1,962 | 2,112 |
| GC (%) | 43.14 | 43.14 |
| Estimated chimera rate in read pairs (%) | 0.03 | 0.03 |
| Genomic read coverage (x) | 42.9 | 42.6 |

| Assembly quality assessment | | |
|---|---|---|
| Genome Scope | | |
| Heterozygosity (%) | 0.16 | 0.22 |
| Read error rate (%) | 0.04 | 0.04 |
| BUSCO | | |
| Completeness (%) | 86.0 | 86.0 |
| Fragmented (%) | 9.7 | 9.7 |
| Lost (%) | 4.3 | 4.3 |
| Bowtie2 overall alignment rate (%) | 98.8 | 98.8 |

**Supplementary Table 3.** Statistical summary of assembly from male-specific reads in *Arapaima gigas.*

| Reads features | |
|---|---|
| Reads no mapped into the female genome | |
| Forward (%) | 2.33 |
| Reverse (%) | 5.50 |
| Unique reads | 897,263 |
| Duplicated reads | 83,571 |
| Average Phred value | 38 |
| GC (%) | 45 |
| Average length (bp) | 176 |

| Assembly features | |
|---|---|
| Total length of bases (Mb) | 2.08 |
| Scaffold N50 (bp) | 3,542 |
| Total number of contigs | 903 |
| Contig N50 (bp) | 3,417 |
| GC (%) | 43.59 |
| Estimated chimera rate in read pairs (%) | 0.02 |
| Genomic read coverage (x) | 93.1 |
| N's per 100 kbp | 9.82 |
| Unmapped female reads (%) | 98 |

| Assembly quality assessment | |
|---|---|
| Genome Scope | |
| Heterozygosity (%) | 1.74 |
| Read error rate (%) | 1.67 |
| Genome haploid length (Mb) | 3.09 |
| Genome unique length (Mb) | 2.98 |
| Genome repeat length (bp) | 102.20 |
| Model fit (%) | 97 |
| BUSCO completeness (%) | 0.4 |

**Supplementary Table 4**. Genes of interest[(1)] and housekeeping genes[(2)], efficiency (E) and $R^2$ values from duplex qPCR assays.

| Gene | Cellular function | Sequence (5' ---> 3') | Sex | Tm (°C) | Amplicon size (bp) | (E) | $R^2$ |
|---|---|---|---|---|---|---|---|
| *MSR_107*[1] | Not available | Not shown | Male | 62 | 107 | 1.09 | 0.994 |
| *MSR_129*[1] | Not available | Not shown | Male | 62 | 129 | 1.01 | 0.998 |
| *TBPL1*[2] | Mediates the transcription of most ribosomal proteins through specialized system | F: ctctggcttgtagatgacattgg | Male | 62 | 205 | 0.96 | 0.991 |
| | | R: tttgttgcgtgtctctgtgg | Female | 62 | 205 | 0.98 | 0.994 |
| *RPL13*[2] | Protein synthesis | F: gggaacaggatgagtttggag | Male | 62 | 237 | 0.95 | 0.996 |
| | | R: gaaagcaagcgtgtgaatgtg | Female | 62 | 237 | 0.97 | 0.993 |
| *EF1A1*[2] | Enzymatic delivery of aminoacyl tRNAs to the ribosome | F: taatgggtgttgccgtttcac | Male | 62 | 251 | 0.94 | 0.995 |
| | | R: tcaaagcctcaaacagaccaag | Female | 62 | 251 | 0.96 | 0.998 |
| *PGK1*[2] | Catalyzes one of the two ATP producing in the glycolytic pathway | F: ggcttgtgaacctcattctgtg | Male | 62 | 233 | 1.02 | 0.994 |
| | | R: aatgcagggcactttgctc | Female | 62 | 233 | 1.04 | 0.996 |

**Supplementary Table 5:** Raw data of duplex qPCR assays for each individual tested in this study (UDT: Undetected; NA: Not available).

MSR_107

| ID | Gonad | | | Fin | | | Mucus | | |
|---|---|---|---|---|---|---|---|---|---|
| | Cq value | Tm (°C) of HKG | Tm (°C) of MSR | Cq value | Tm (°C) of HKG | Tm (°C) of MSR | Cq value | Tm (°C) of HKG | Tm (°C) of MSR |
| Female_1 | 23,74 | 84,68 | UDT | NA | NA | NA | NA | NA | NA |
| Female_2 | 24,28 | 84,64 | UDT | NA | NA | NA | NA | NA | NA |
| Female_3 | 23,73 | 84,68 | UDT | NA | NA | NA | NA | NA | NA |
| Female_4 | 22,68 | 84,65 | UDT | NA | NA | NA | NA | NA | NA |
| Female_5 | 22,22 | 84,58 | UDT | NA | NA | NA | NA | NA | NA |
| Female_6 | 22,53 | 84,62 | UDT | NA | NA | NA | NA | NA | NA |
| Female_7 | 22,48 | 84,51 | UDT | NA | NA | NA | NA | NA | NA |
| Female_8 | 22,64 | 84,51 | UDT | NA | NA | NA | NA | NA | NA |
| Female_9 | 22,89 | 84,53 | UDT | NA | NA | NA | NA | NA | NA |
| Female_10 | 22,37 | 84,61 | UDT | 24,05 | 84,87 | UDT | 23,41 | 84,90 | UDT |
| Female_11 | 23,49 | 84,55 | UDT | 24,33 | 84,65 | UDT | 23,44 | 84,70 | UDT |
| Female_12 | 22,55 | 84,55 | UDT | 24,30 | 84,64 | UDT | 23,18 | 84,71 | UDT |
| Female_13 | 21,99 | 84,71 | UDT | 24,96 | 84,91 | UDT | 23,17 | 84,88 | UDT |
| Female_14 | 22,04 | 84,54 | UDT | 24,71 | 84,64 | UDT | 23,52 | 84,71 | UDT |
| Female_15 | 22,62 | 84,68 | UDT | 24,26 | 84,88 | UDT | 23,65 | 84,91 | UDT |
| Female_16 | 21,41 | 84,72 | UDT | 24,63 | 84,82 | UDT | 23,58 | 84,87 | UDT |
| Female_17 | 22,68 | 84,69 | UDT | 23,09 | 84,89 | UDT | 24,31 | 84,99 | UDT |
| Female_18 | 23,09 | 84,45 | UDT | 23,39 | 84,95 | UDT | 23,49 | 84,77 | UDT |
| Female_19 | 22,78 | 84,52 | UDT | 22,81 | 84,73 | UDT | 25,00 | 84,59 | UDT |
| Female_20 | 23,16 | 84,52 | UDT | 21,38 | 84,91 | UDT | 25,00 | 84,71 | UDT |
| Female_21 | 22,97 | 84,75 | UDT | 23,63 | 84,99 | UDT | 23,04 | 84,75 | UDT |
| Female_22 | 22,67 | 84,46 | UDT | 23,00 | 84,72 | UDT | 24,24 | 84,62 | UDT |
| Female_23 | 22,81 | 84,48 | UDT | 23,23 | 84,82 | UDT | 23,57 | 84,74 | UDT |
| Female_24 | 22,62 | 84,46 | UDT | 23,07 | 84,70 | UDT | 24,03 | 84,58 | UDT |
| Female_25 | 23,15 | 84,77 | UDT | 23,56 | 84,70 | UDT | 23,59 | 84,57 | UDT |
| Female_26 | 22,83 | 84,44 | UDT | NA | NA | NA | NA | NA | NA |
| Female_27 | 22,81 | 84,51 | UDT | NA | NA | NA | NA | NA | NA |
| Female_28 | 22,99 | 84,76 | UDT | NA | NA | NA | NA | NA | NA |
| Male_1 | 23,20 | 84,82 | 77,64 | NA | NA | NA | NA | NA | NA |
| Male_2 | 23,23 | 84,81 | 76,67 | NA | NA | NA | NA | NA | NA |
| Male_3 | 22,41 | 84,85 | 76,62 | NA | NA | NA | NA | NA | NA |
| Male_4 | 20,98 | 84,56 | 76,46 | NA | NA | NA | NA | NA | NA |
| Male_5 | 20,87 | 84,59 | 76,47 | NA | NA | NA | NA | NA | NA |
| Male_6 | 20,95 | 84,55 | 76,55 | NA | NA | NA | NA | NA | NA |
| Male_7 | 20,89 | 84,56 | 76,53 | NA | NA | NA | NA | NA | NA |
| Male_8 | 21,02 | 84,64 | 76,53 | NA | NA | NA | NA | NA | NA |
| Male_9 | 21,15 | 84,61 | 76,51 | NA | NA | NA | NA | NA | NA |
| Male_10 | 20,99 | 84,66 | 76,44 | NA | NA | NA | NA | NA | NA |
| Male_11 | 21,38 | 84,69 | 76,67 | NA | NA | NA | NA | NA | NA |
| Male_12 | 20,90 | 84,65 | 76,48 | NA | NA | NA | NA | NA | NA |
| Male_13 | 20,49 | 84,89 | 76,59 | 21,80 | 84,84 | 76,55 | 21,22 | 84,88 | 76,62 |
| Male_14 | 20,66 | 84,77 | 76,72 | 21,57 | 84,75 | 76,68 | 21,46 | 84,70 | 76,64 |
| Male_15 | 20,53 | 84,69 | 76,65 | 21,23 | 84,60 | 76,51 | 21,74 | 84,76 | 76,66 |
| Male_16 | 20,88 | 84,99 | 76,61 | 21,14 | 84,88 | 76,58 | 21,53 | 84,87 | 76,60 |
| Male_17 | 20,59 | 84,54 | 76,50 | 21,20 | 84,69 | 76,61 | 21,49 | 84,69 | 76,61 |
| Male_18 | 20,59 | 84,74 | 76,41 | 21,06 | 84,78 | 76,52 | 21,46 | 84,83 | 76,60 |
| Male_19 | 20,63 | 84,51 | 76,50 | 20,87 | 84,66 | 76,60 | 21,47 | 84,66 | 76,63 |
| Male_20 | 20,64 | 84,55 | 76,53 | 21,09 | 84,68 | 76,61 | 21,50 | 84,67 | 76,59 |
| Male_21 | 20,92 | 84,48 | 76,46 | 20,68 | 84,80 | 76,69 | 21,61 | 84,45 | 76,48 |
| Male_22 | 22,73 | 84,93 | 76,59 | 20,74 | 84,75 | 76,80 | 21,90 | 84,77 | 76,65 |
| Male_23 | 21,16 | 84,90 | 76,58 | 20,85 | 85,06 | 76,73 | 21,59 | 84,83 | 76,57 |
| Male_24 | 21,88 | 84,52 | 76,52 | 21,06 | 84,75 | 76,74 | 21,41 | 84,59 | 76,52 |
| Male_25 | 20,83 | 84,86 | 76,60 | NA | NA | NA | NA | NA | NA |
| Male_26 | 21,00 | 84,54 | 76,49 | NA | NA | NA | NA | NA | NA |
| Male_27 | 21,01 | 84,46 | 76,48 | NA | NA | NA | NA | NA | NA |

MSR_129

| ID | Gonad | | | Fin | | | Mucus | | |
|---|---|---|---|---|---|---|---|---|---|
| | Cq value | Tm (°C) of HKG | Tm (°C) of MSR | Cq value | Tm (°C) of HKG | Tm (°C) of MSR | Cq value | Tm (°C) of HKG | Tm (°C) of MSR |
| Female_1 | 23,79 | 84,61 | UDT | NA | NA | NA | NA | NA | NA |
| Female_2 | 23,98 | 84,59 | UDT | NA | NA | NA | NA | NA | NA |
| Female_3 | 24,25 | 84,61 | UDT | NA | NA | NA | NA | NA | NA |
| Female_4 | 22,84 | 84,63 | UDT | NA | NA | NA | NA | NA | NA |
| Female_5 | 22,27 | 84,58 | UDT | NA | NA | NA | NA | NA | NA |
| Female_6 | 22,57 | 84,60 | UDT | NA | NA | NA | NA | NA | NA |
| Female_7 | 22,54 | 84,51 | UDT | NA | NA | NA | NA | NA | NA |
| Female_8 | 22,67 | 84,54 | UDT | NA | NA | NA | NA | NA | NA |
| Female_9 | 22,96 | 84,53 | UDT | NA | NA | NA | NA | NA | NA |
| Female_10 | 22,18 | 84,75 | UDT | 24,23 | 84,87 | UDT | 23,40 | 84,91 | UDT |
| Female_11 | 22,80 | 84,62 | UDT | 24,68 | 84,64 | UDT | 23,40 | 84,71 | UDT |
| Female_12 | 22,54 | 84,49 | UDT | 24,57 | 84,64 | UDT | 23,25 | 84,71 | UDT |
| Female_13 | 22,24 | 84,77 | UDT | 25,06 | 84,93 | UDT | 23,17 | 84,93 | UDT |
| Female_14 | 22,20 | 84,53 | UDT | 24,79 | 84,74 | UDT | 23,59 | 84,71 | UDT |
| Female_15 | 22,67 | 84,75 | UDT | 24,42 | 84,94 | UDT | 23,28 | 84,94 | UDT |
| Female_16 | 22,68 | 84,87 | UDT | 24,87 | 84,95 | UDT | 23,71 | 84,95 | UDT |
| Female_17 | 22,71 | 84,75 | UDT | 23,15 | 84,89 | UDT | 24,38 | 84,92 | UDT |
| Female_18 | 23,15 | 84,60 | UDT | 23,37 | 84,83 | UDT | 23,59 | 84,91 | UDT |
| Female_19 | 22,79 | 84,52 | UDT | 22,96 | 84,70 | UDT | 23,38 | 84,62 | UDT |
| Female_20 | 23,21 | 84,59 | UDT | 23,07 | 84,92 | UDT | 24,25 | 84,72 | UDT |
| Female_21 | 23,13 | 84,90 | UDT | 23,52 | 84,90 | UDT | 23,27 | 84,82 | UDT |
| Female_22 | 22,81 | 84,48 | UDT | 23,03 | 84,69 | UDT | 23,41 | 84,65 | UDT |
| Female_23 | 22,85 | 84,46 | UDT | 23,42 | 84,88 | UDT | 23,75 | 84,82 | UDT |
| Female_24 | 22,61 | 84,47 | UDT | 23,15 | 84,73 | UDT | 23,81 | 84,59 | UDT |
| Female_25 | 23,31 | 84,94 | UDT | 23,59 | 84,69 | UDT | 24,13 | 84,62 | UDT |
| Female_26 | 22,75 | 84,42 | UDT | NA | NA | NA | NA | NA | NA |
| Female_27 | 23,01 | 84,73 | UDT | NA | NA | NA | NA | NA | NA |
| Female_28 | 23,13 | 84,91 | UDT | NA | NA | NA | NA | NA | NA |
| Male_1 | 23,19 | 84,80 | 77,52 | NA | NA | NA | NA | NA | NA |
| Male_2 | 23,16 | 84,95 | 77,49 | NA | NA | NA | NA | NA | NA |
| Male_3 | 22,25 | 84,95 | 77,49 | NA | NA | NA | NA | NA | NA |
| Male_4 | 20,80 | 84,60 | 77,30 | NA | NA | NA | NA | NA | NA |
| Male_5 | 20,99 | 84,59 | 77,26 | NA | NA | NA | NA | NA | NA |
| Male_6 | 20,90 | 84,58 | 77,32 | NA | NA | NA | NA | NA | NA |
| Male_7 | 20,82 | 84,59 | 77,34 | NA | NA | NA | NA | NA | NA |
| Male_8 | 21,09 | 84,60 | 77,30 | NA | NA | NA | NA | NA | NA |
| Male_9 | 20,96 | 84,69 | 77,41 | NA | NA | NA | NA | NA | NA |
| Male_10 | 21,17 | 84,62 | 77,37 | NA | NA | NA | NA | NA | NA |
| Male_11 | 20,89 | 84,76 | 77,37 | NA | NA | NA | NA | NA | NA |
| Male_12 | 20,26 | 84,89 | 77,36 | NA | NA | NA | NA | NA | NA |
| Male_13 | 20,83 | 84,70 | 77,42 | 21,15 | 85,00 | 77,51 | 21,17 | 85,07 | 77,58 |
| Male_14 | 20,54 | 84,89 | 77,65 | 21,51 | 84,92 | 77,62 | 21,31 | 84,87 | 77,61 |
| Male_15 | 20,42 | 84,71 | 77,47 | 21,05 | 84,83 | 77,53 | 21,80 | 85,00 | 77,65 |
| Male_16 | 20,74 | 84,93 | 77,54 | 21,12 | 85,06 | 77,58 | 21,42 | 85,03 | 77,57 |
| Male_17 | 20,50 | 84,89 | 77,65 | 21,18 | 84,91 | 77,60 | 21,40 | 84,86 | 77,56 |
| Male_18 | 20,58 | 84,99 | 77,43 | 21,01 | 85,02 | 77,51 | 21,44 | 85,04 | 77,61 |
| Male_19 | 20,60 | 84,79 | 77,55 | 20,89 | 84,93 | 77,67 | 21,44 | 84,86 | 77,59 |
| Male_20 | 20,56 | 84,89 | 77,65 | 21,08 | 84,92 | 77,59 | 21,46 | 84,90 | 77,58 |
| Male_21 | 20,82 | 84,73 | 77,48 | 20,44 | 84,87 | 77,54 | 22,01 | 84,68 | 77,25 |
| Male_22 | 22,64 | 84,92 | 77,40 | 20,85 | 85,07 | 77,68 | 21,84 | 85,02 | 77,53 |
| Male_23 | 21,04 | 84,95 | 77,49 | 20,64 | 85,10 | 77,53 | 20,76 | 85,02 | 77,31 |
| Male_24 | 21,85 | 84,75 | 77,51 | 20,82 | 84,70 | 77,51 | 20,70 | 84,58 | 77,42 |
| Male_25 | 20,87 | 84,92 | 77,39 | NA | NA | NA | NA | NA | NA |
| Male_26 | 20,89 | 84,80 | 77,52 | NA | NA | NA | NA | NA | NA |
| Male_27 | 20,89 | 84,72 | 77,51 | NA | NA | NA | NA | NA | NA |
